# Sex-Specific Cardiovascular Adaptations to Simulated Microgravity in Sprague-Dawley Rats

**DOI:** 10.1101/2024.03.29.587264

**Authors:** Ebrahim Elsangeedy, Dina N. Yamaleyeva, Nicholas P. Edenhoffer, Allyson Deak, Anna Soloshenko, Jonathan Ray, Xuming Sun, Omar H. Shaltout, Nildris Cruz Diaz, Brian Westwood, Daniel Kim-Shapiro, Debra I. Diz, Shay Soker, Victor M. Pulgar, April Ronca, Jeffrey S. Willey, Liliya M. Yamaleyeva

## Abstract

Men and women have different cardiovascular responses to spaceflight; however few studies have focused on direct comparisons between sexes. Therefore, we investigated cardiovascular system differences, including arterial stiffness between socially and sexually mature 20-week-old male and female Sprague Dawley (SD) rats exposed to hindlimb unloading (HLU) - an analogue for spaceflight-induced microgravity. Two weeks of HLU had no effect on body weight in either male or female rats. The index of arterial stiffness determined by ultrasound, pulse wave velocity (PWV), was greater in the aortic arch and carotid artery of females after HLU versus control females. HLU had no effect on arterial PWV in males. α smooth muscle actin, myosin, collagen, elastin, and collagen-to-elastin ratio were not different in rats of either sex in response to HLU. HLU exposure did not alter individual collagen fiber characteristics in studied groups. The levels of G protein-coupled estrogen receptor (GPER) were lower in the aorta of SD females exposed to HLU compared with female controls but not in males. These changes were associated with lower PPAR γ and increased oxidative stress markers (8-hydroxy-2’-deoxyguanosine and p47phox) in the females. Diastolic cardiac function was altered in females after HLU versus control females. GPER agonist, G1 prevented the increase in pulse wave velocity and 8-hydroxy-2’-deoxyguanosine, without altering PPAR γ or p47phox. Our data revealed that lower GPER in the HLU females contributes to the development of arterial stiffness, and that the SD rat is a suitable model to study the cardiovascular response of females to HLU.

## 1 Introduction

The cardiovascular adaptations in astronauts both during and after spaceflight are variable in terms of sex, duration of exposure, and the model used for the studies. Men and women have different recovery mechanisms from stress factors, which can influence the response to the spaceflight environment (1). Though women are known to be protected against heart disease prior to menopause, responses of the vascular system to spaceflight are complicated by additional stress factors including microgravity, space radiation, inactivity, or isolation. Few studies have focused on the sex differences in the adaptations of the cardiovascular system to spaceflight in spite of the fact that the proportion of women participating in space missions has reached about 10% and continues to increase (2). The empirically reported effects of spaceflight on the cardiovascular system are varied across people who have been in space (3). Some common short-term physiological effects of spaceflight include changes in blood volume distribution which cause orthostatic intolerance and augmented baroreflex response (3). Prolonged exposure to spaceflight impacted red blood cell count and increased cardiac output at six months post-flight (4, 5). Sex-based differences in cardiovascular responses have been reported with changes in both sexes occurring after several weeks in space (1, 6). Female astronauts experience a greater loss of plasma volume while males develop a spaceflight associated neuro-ocular syndrome (SANS) (7–9). Furthermore, after a 6-month spaceflight, women had increased renin levels, while both men and women experienced increased carotid artery stiffness (10). The observed sex differences underscore the importance of understanding the underlying mechanisms of cardiovascular changes and sex-specific adaptations to both terrestrial and space-induced stressors important for the development of personalized preventative countermeasures for astronauts. To better understand the changes in cardiovascular adaptations in response to spaceflight environment, terrestrial studies are focused on recapitulating microgravity in rodent models with hindlimb unloading (HLU) (11). However, the majority of these studies use male rodent subjects, with even fewer examining differences between sexes. The mechanisms underlying the cardiovascular system responses and arterial stiffness are not well understood particularly in the female subjects. Therefore, the present study investigated the mechanisms underlying the development of central arterial stiffness in female and male Sprague Dawley (SD) rats in response to HLU as an analogue for spaceflight-induced microgravity.

## 2 Materials and Methods

### 2.1 Animals

The study was approved by the Institutional Animal Care and Use Committee of the Wake Forest University School of Medicine (A21-039). Female and male Sprague-Dawley (SD) rats were purchased from Charles River Laboratories (Wilmington, MA, USA). All rats were housed at 12:12-h light-dark cycle, a constant room temperature and humidity. A standard rodent chow (Lab Diet 5P00 - Prolab RMH 3000, PMI Nutrition International, INC, Brentwood, MO) and water were given *ad libitum* throughout the experimental protocols.

### 2.2 Study timeline

Socially and sexually mature female and male SD rats at 20 weeks of age were randomly assigned to the following groups: female control (F), female HLU (F HLU), male control (M), or male HLU (M HLU). HLU groups underwent hindlimb unloading for 14-days (12). Control groups were exposed to the same cages for a 14-day period. On the morning of day 14, all groups were imaged with ultrasound followed by EchoMRI, and then sacrificed to collect tissues. Additional groups of female rats underwent treatment with G1, a G protein-coupled estrogen receptor (GPER) agonist, delivered via osmotic minipumps for three weeks. Pumps were inserted one-week prior to HLU exposure and continued for two weeks during the HLU exposure. Aortic pulse wave velocity (PWV) and injury markers were analyzed at the end of the 3-week treatment with G1.

### 2.3 Hindlimb unloading (HLU)

To determine the effects of microgravity, 20-week-old male and female SD rats were suspended by the tail via HLU for 14 days resulting in a head-down body angle of 30-35 degrees. Age-matched, non-suspended controls remained in similar cages (12). Animals were monitored for signs of distress by monitoring food and water intake and loss of body weight throughout the exposure to HLU.

### 2.4 Ultrasound assessment of arterial stiffness

Rats were anesthetized with isoflurane (1.5%) and placed on a temperature-controlled platform. Electrocardiogram (ECG) and heart rate were monitored and recorded throughout the imaging protocol. Hair was removed from the thoracic and neck areas of rats with a depilatory cream prior to imaging. Pulse wave velocity (PWV) of the aortic arch and common carotid arteries were determined in male and female rats using a MS250S transducer and high-resolution ultrasound imaging system Vevo 2100 LAZR (FujiFilm, VisualSonics). The PWV of the aortic arch and carotid arteries was measured by the following equation: the D (mm), distance from ascending to descending artery, divided by time interval, Δt, which represent the difference between t2 and t1 (ms). Time intervals between the R-wave of the ECG to the foot of the Doppler waveforms were averaged over at least three cardiac cycles, and the pulse-transit time was calculated by subtracting the mean ascending aortic arch (t1) time interval from the mean descending aortic arch (t2) time interval.

### 2.5 Body composition

EchoMRI^TM^ Analyzer was used to determine the body composition in studied groups.

### 2.6 Collagen content

Aortas were fixed in formalin and ethanol followed by the paraffin-embedding. Aortic sections were cut at thickness of 5 µm. Collagen content was determined by Masson’s trichrome staining (Abcam ab150686). Using a Hamamatsu Nanozoomer 2.0 HT equipped with NDP.scan and NDP.view, slides containing stained aortic sections were imaged at 20X magnification. Using Adobe Photoshop 7.0, each slide was quantified for the total area of collagen deposition (red, orange, yellow, or green) and reported as a collagen-to-tissue ratio.

### 2.7 Collagen fiber properties

5 µm sections were stained with Picrosirius red (Sigma Aldrich, St. Louis, MO; assay components Cat#: 197378-100G; P6744-16A; 36-554-8) to assess collagen fiber properties. Slides were imaged using polarized light, and 90 x 90 µm representative regions of interest (ROIs) were selected around aortic arch to determine collagen fiber bundling, length, width, straightness, and alignment. Collagen bundling was assessed using an R script that binned hue values in each ROI into red, orange, yellow, and green, in order of high to low collagen bundling (13, 14). Data was presented as a ratio of the binned values to the total number of pixels binned.

Collagen fiber length, width, and straightness were determined through CT-FIRE software (Laboratory for Optical and Computational Instrumentation, The University of Wisconsin-Madison) using default settings (15). Coefficient of alignment was determined through CurveAlign software (Laboratory for Optical and Computational Instrumentation, The University of Wisconsin-Madison), using default settings (15).

#### Smooth muscle fiber characterization

Extracellular matrix (ECM) characteristics were analyzed using MATLAB and the CT-FIRE algorithm to assess fiber straightness, fiber length, and fiber width. Fiber angle was quantified at the individual fiber level. CT-FIRE excludes empty pixels prior to analysis. Quantification of fibers was done using CT-FIRE and Curve Align scripts (14).

### 2.8 Elastin fiber content

*Verhoeff’s stain* (American MasterTech; Cat#: KTVEL) was used to quantify elastin content in aortic sections using the manufacturer’s protocol. The staining intensity in the tunica media layer of the aorta was normalized to the intensity of the background and to the maximal value of 255 of the red, green, and blue intensity (RGB) component and reported as relative intensity units (16).

### 2.9 Immunohistochemistry for morphological, inflammatory, and oxidative stress markers

Tissues were fixed in 10% formalin followed by 70% ethanol, embedded in paraffin, and cut into 5 µm sections. Immunostaining was performed using the avidin biotin complex (ABC) method with 0.1% diaminobenzene solution used as the chromogen as previously described (15). Antigen retrieval treatment with sodium citrate buffer (pH 6.0) was applied at 90-95°C for 30 min. Non-specific binding was blocked in a buffer containing 10% normal goat serum, 0.1% Triton-X in PBS for 30 min. Aortic sections were incubated with goat anti-*α-*smooth muscle actin (1:500 dilution; Novus Biologicals, Centennial, CO, USA; Cat#: NB300-978), rabbit anti-smooth muscle myosin heavy chain 11 (1:500 dilution; Abcam, Waltham, MA, USA; Cat#: ab53219), rabbit anti-4-hydroxynonenal (4-HNE; 1:10,000 dilution; Abcam, Waltham, MA, USA; Cat#: ab46545), rabbit anti-p47 phox (1:400 dilution; LifeSpan BioSciences, Shirley, MA, USA; Cat#: LS-C331296), mouse anti-8-hydroxy-2’-deoxyguanosine (8-OHdG; 1:100 dilution; JaICA, Fukuroi, Shizuoka, Japan; Cat#: MOG-020P), rabbit anti-cyclooxygenase-2 (COX-2, 1:200 dilution; Cayman Chemicals, Ann Arbor, MI, USA; Cat#: 160126), mouse anti-endothelial nitric oxide synthase (eNOS, 1:100 dilution; BD Biosciences, Franklin Lakes, NJ; Cat#: 610297), mouse anti-macrophages/monocytes antibody, clone ED-1/CD68 (1:200 dilution; MilliporeSigma, Burlington, MA, USA; Cat#: MAB1435), rabbit anti-peroxisome proliferator-activated receptor gamma (PPARγ; 1:1200 dilution; Biorbyt, LLC, Durham, NC, USA; Cat#: orb11291), rabbit anti-GPER (1:200 dilution; Biorbyt, LLC, Durham, NC, USA; Cat#: orb10740) and secondary biotinylated anti-rabbit, mouse or goat antibodies (1:500 dilution; Vector Laboratories, Burlingame, CA, USA). Slides were scanned with a Hamamatsu Nanozoomer 2.0 HT utilizing NDP.scan and NDP.view as the imaging software at 20X magnification. Images were analyzed in four fields per aortic cross-section using Adobe Photoshop 7.0 by the investigator blinded to the experimental groups. The staining was normalized to the intensity of the background and to the maximal value of the RGB component and reported as relative intensity units as described (16).

### 2.10 Statistics

Groups were compared using one-way or two-way analysis of variance (ANOVA) followed by the Bonferroni post-hoc tests (GraphPad Software, San Diego, CA). A p-value less than 0.05 was considered statistically significantly different. All data were presented as mean ± SEM.

## 3 Results

### 3.1 Effect of HLU on body and organ weight

The body weight of male control rats was significantly (p<0.05) greater than the body weight of female control rats (**Figure 1A**). Similarly, male HLU rats were heavier than female HLU rats (**Figure 1A**). To account for this variability in body weight while allowing for comparison between sexes, organ weights (heart, kidney, pancreas, spleen, uterus) were normalized to individual body weight. Body composition analysis revealed no differences in total fat mass between study groups, however males had greater lean mass (**Figure 1B-C**). Exposure to HLU reduced lean mass in males (Figure 1C). Within each sex, HLU treatment did not affect organ weights (**Figure 1D, F, G**) except the kidneys where female rats exposed to HLU had increased kidney weight compared to the female control group (**Figure 1E**). HLU did not affect uterine weight in female SD rats (**Figure 1H**).

**Figure 1.**
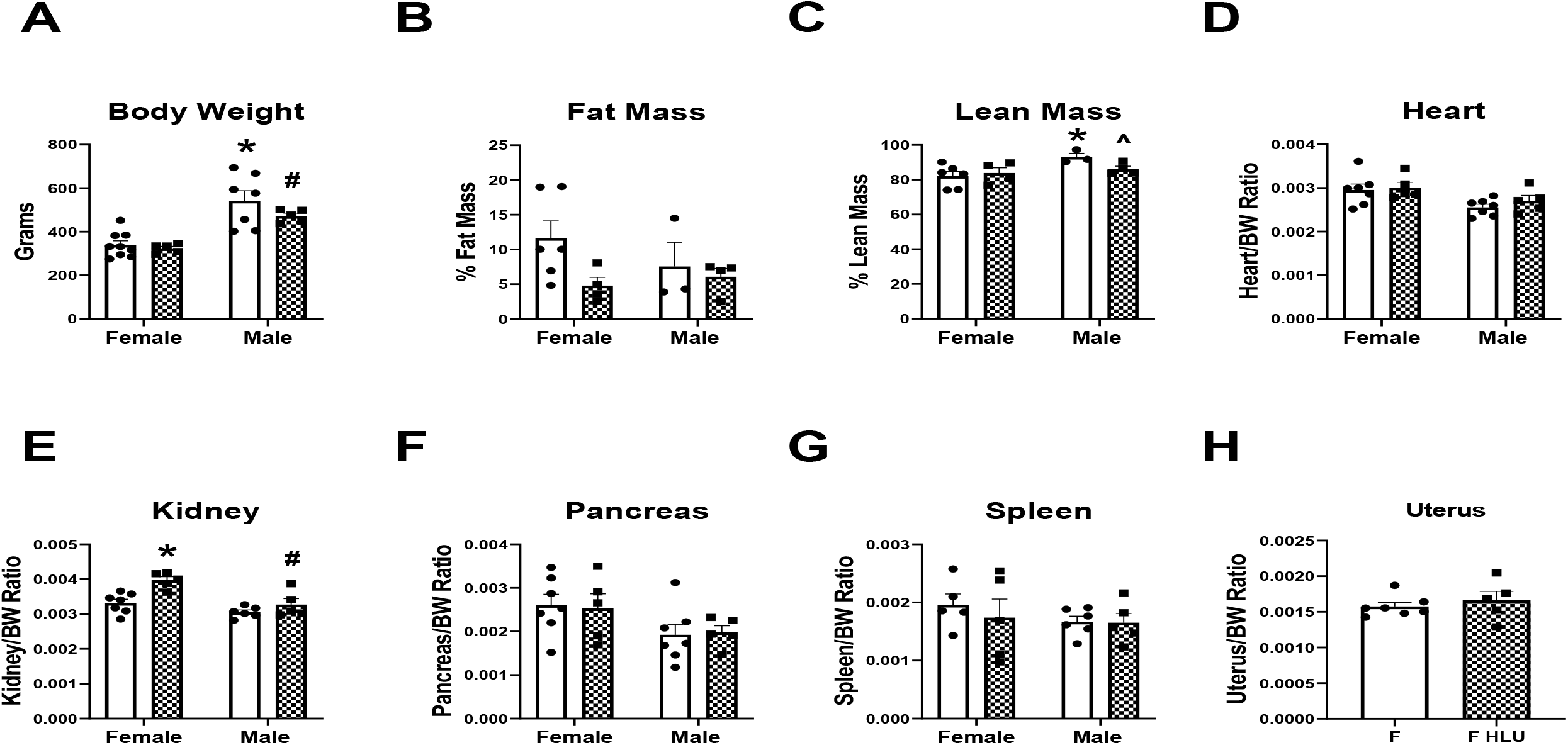
The effect of 14-days of HLU on body and organ weights in 20-week-old female and male Sprague-Dawley rats. Clear bars indicate control unexposed animals; checkered bars indicate rats exposed to hindlimb unloading (HLU). Data are mean ± SEM, *p<0.05 vs. Control Female; ^#^p<0.05 vs. HLU Female; ^^^p<0.05 vs. Control Male.

### 3.2 Aortic PWV and cardiac function in response to HLU

Pulse wave velocity (PWV) of the aortic arch in male and female SD rats was determined using the high-resolution ultrasound imaging system Vevo LAZR, (**Figure 2**). HLU females had a greater aortic arch PWV compared to control females or HLU males (**Figure 2**). No differences were found in PWV in HLU males compared with male controls (**Figure 2**). HLU also increased PWV of the left common carotid artery in the female rats compared to female controls **(Figure S1)**. Aortic wall media thickness was not different in male groups after HLU exposure. To determine whether changes in cardiac function could explain differences in PWV caused by HLU exposure, systolic and diastolic function parameters were measured in the female control and HLU rats. Stroke volume and cardiac output were not different between control and HLU groups **(Figure S2 A, B)**. Although within normal ranges, female HLU rats had increased ejection fraction compared to control female rats **(Figure S2C)**. Diastolic function, measured by E/E’ ratio was worse in HLU female rats versus controls **(Figure S2 D)**.

**Figure 2.**
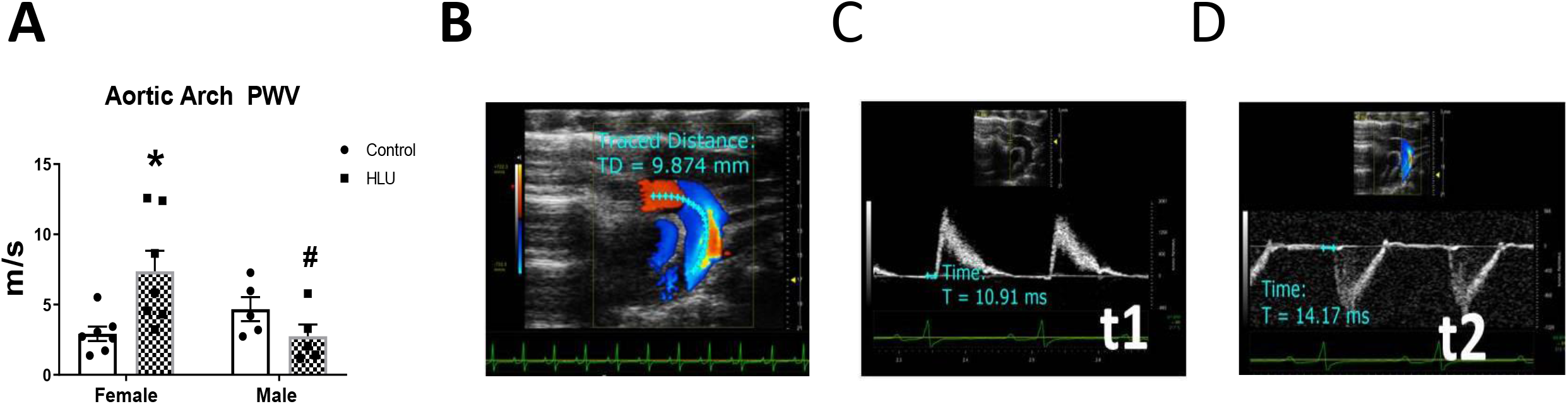
Aortic arch pulse wave velocity (PWV) in female and male rats in response to HLU. Clear bars indicate control unexposed animals; checkered bars indicate rats exposed to hindlimb unloading (HLU) **(A)**. Data are mean ± SEM, *p<0.05 vs. Control Female; ^#^p<0.05 vs. HLU Female. PWV was calculated as PWV=D/pulse transit time Δt, where D is the distance between two points in mm (t2 and t1) in the aortic arch, and t is the signal transit time estimated between R on electrocardiogram and the beginning of the Doppler wave in ms **(B-D).**

### 3.3 Characterization of aortic extracellular matrix (ECM) and contractile components

α-smooth muscle actin and myosin were assessed in the intima of aortic arch of female and male rats following HLU exposure (**Figure 3**). The aortas of male controls had greater levels of both proteins compared to control females. HLU had no effect on either α-smooth muscle actin or myosin in female or male rats (**Figure 3**). Collagen and elastin content, and collagen-to-elastin ratio were not different in control or HLU female or male rats (**Figure 4**).

**Figure 3.**
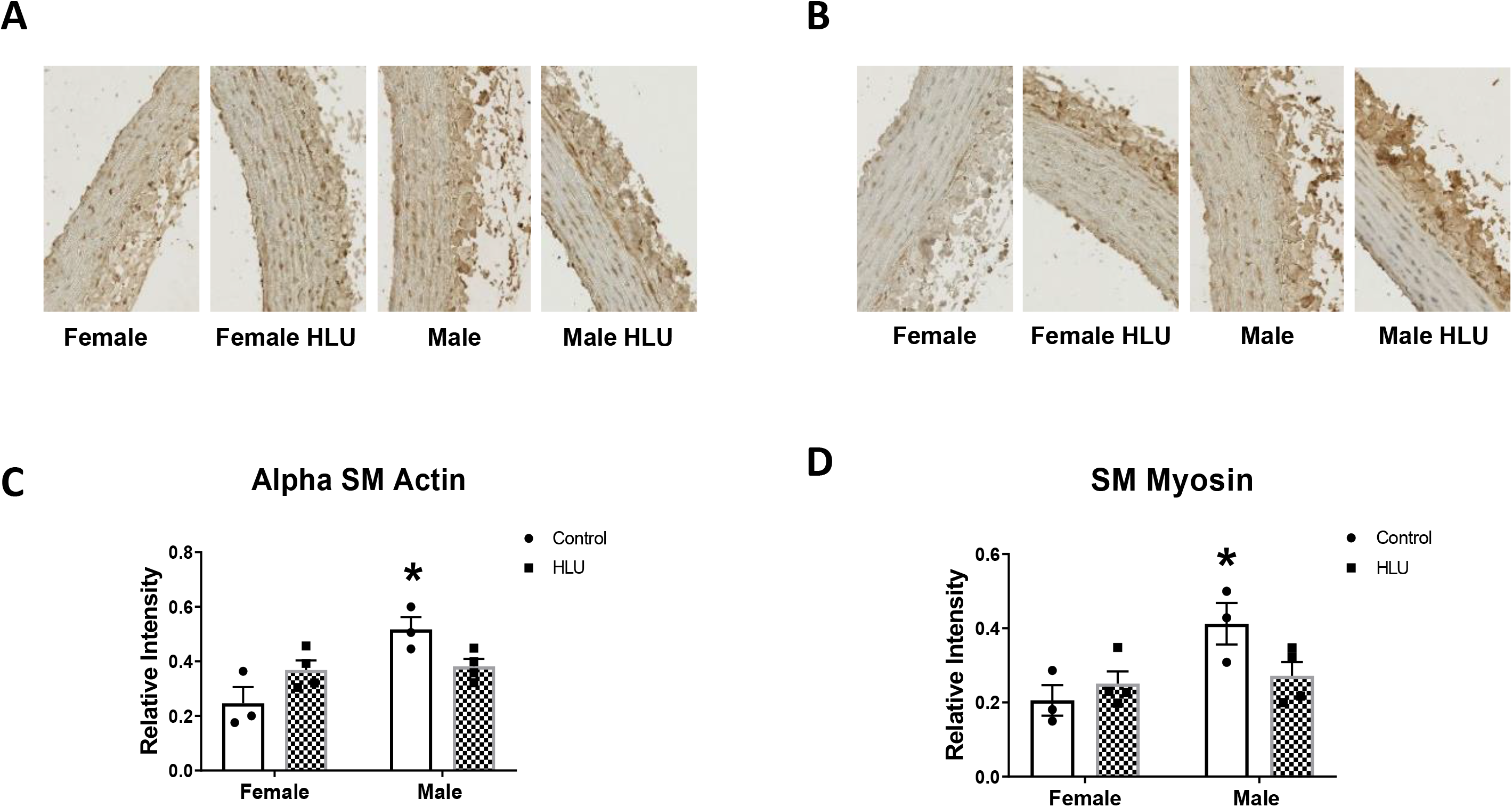
Smooth muscle (SM) contractile protein staining of aortic media. Representative images and analysis of relative intensity of α-smooth muscle alpha actin **(A and C)** and smooth muscle myosin **(B and D)** in female and male SD rats with or without HLU exposure are shown. Data are mean ± SEM, *p<0.05, vs. Female, n=3-4 per group.

**Figure 4.**
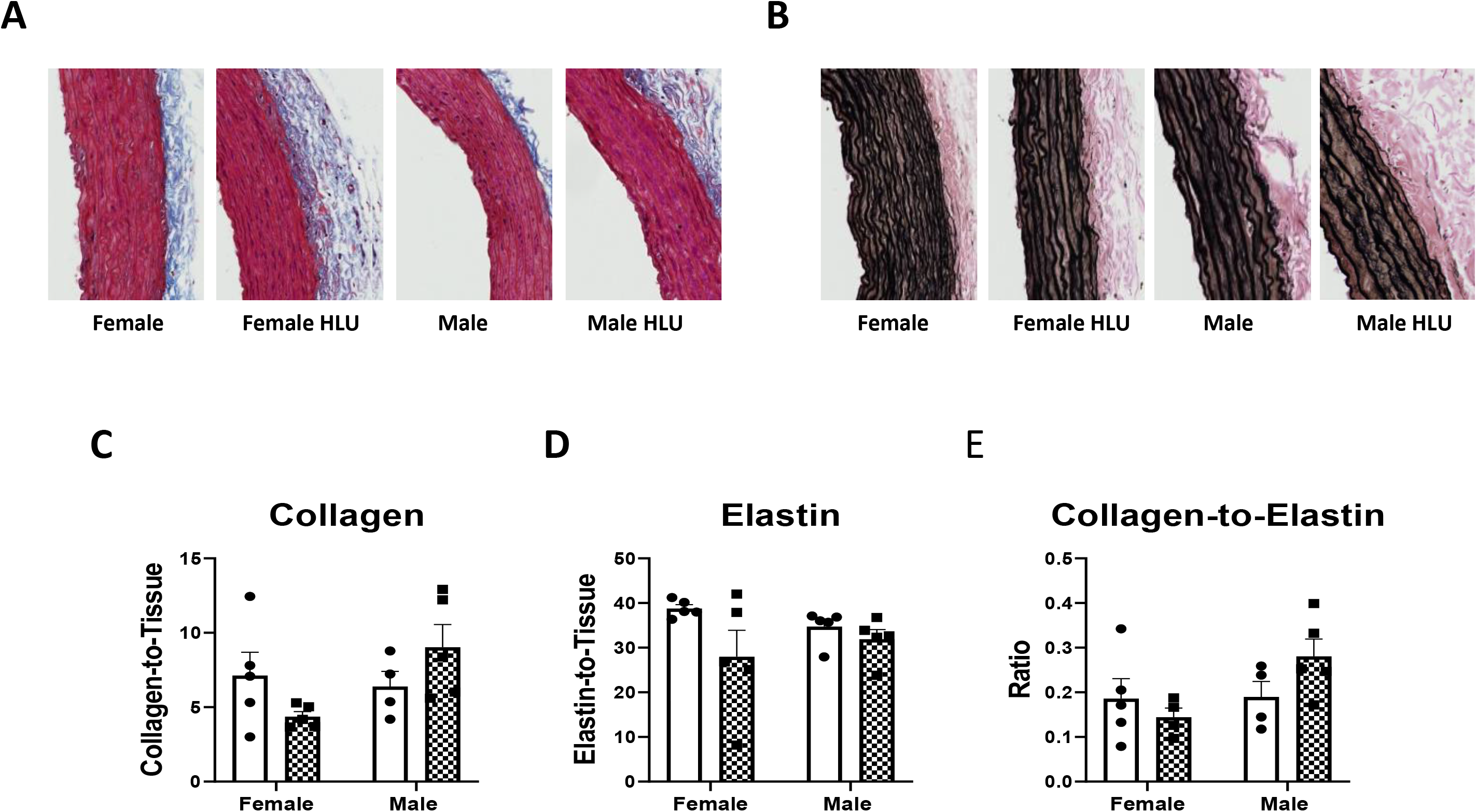
Effect of HLU on total collagen or elastin area, and collagen-to-elastin ratio in aortic media of SD rats. Aortas were stained for collagen by Picrosirius Red stain **(A, C)** and elastin by Verhoeff’s stain **(B, D)**. Data are mean ± SEM, n=4-5.

Collagen bundling in the aorta was measured by hue analysis to gain understanding of the changes in collagen density after HLU and between sexes (**Figure 5**). Results were normalized to the number of total pixels. Through picrosirius red staining and imaging with polarized light, highly dense collagen appears red while very loosely bundled collagen appears green, with orange and yellow colors representing intermediary values (**Figure 5**). Hue analysis determined collagen content based on standardized binning of colors as previously reported (13, 14). All groups followed a similar collagen bundling pattern where prevalence of dense fibers was the highest while the prevalence of fibers with loose collagen was the lowest (**Figure 5**). Female control rats had greater red collagen compared with male controls (**Figure 5**). However, HLU exposure did not influence the dense collagen fibers (red) in female rats in comparison to control females. Less bundled collagen (yellow and green) was lower in female controls compared to male controls (**Figure 5**). HLU did not affect the prevalence of the loosely bundled collagen (yellow or green) in any of the groups. When the data were not normalized to total pixels, female HLU rats had significantly a higher amount of highly dense collagen bundling (red) compared to all other groups **(Figure S3)**.

**Figure 5.**
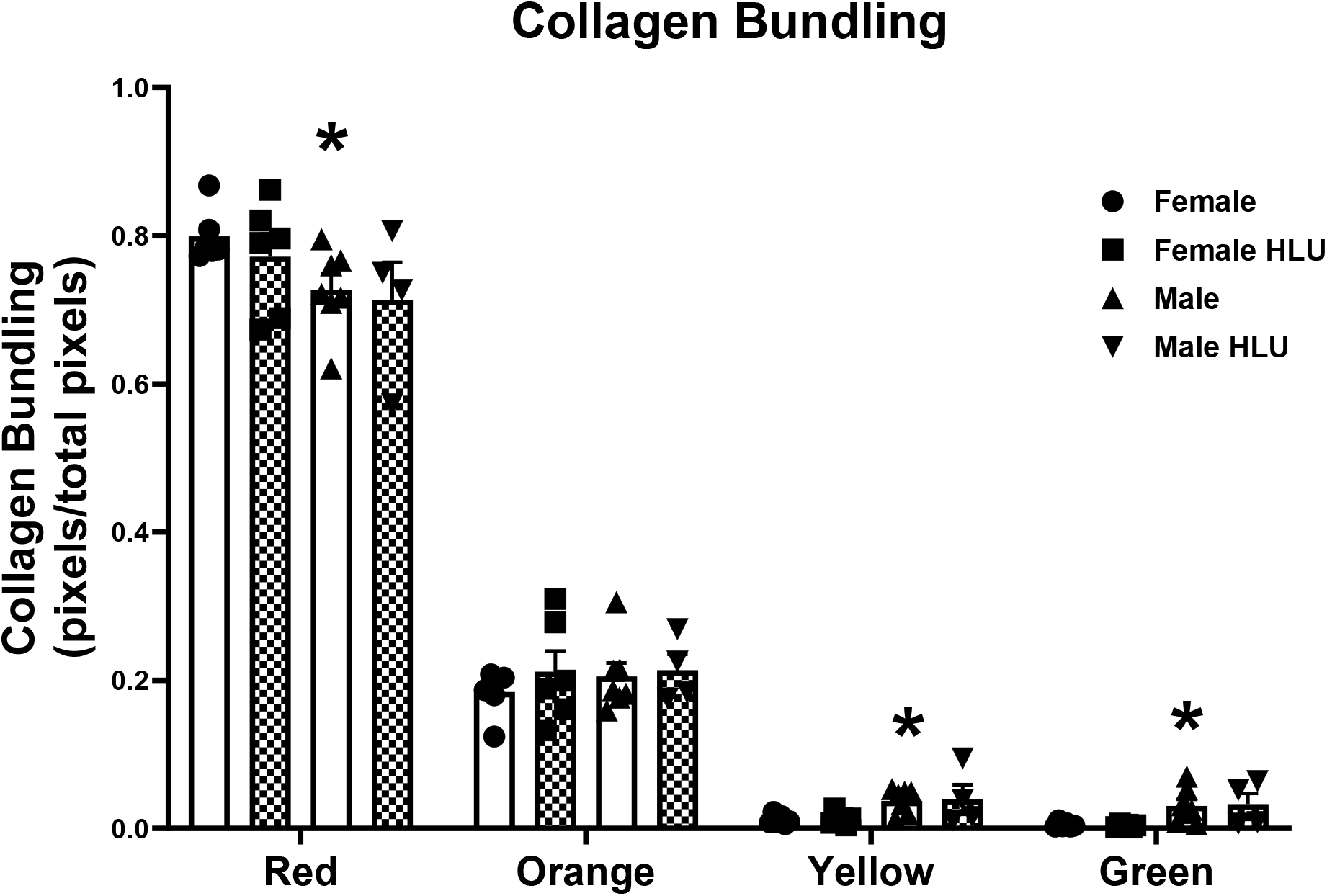
Collagen fiber bundling in the aorta of female and male SD rats in response to HLU. Clear bars indicate control animals; checkered bars indicate rats exposed to hindlimb unloading (HLU). Picrosirius red stained slides were imaged with polarized light, and the pixel values of representative regions were binned into red, orange, yellow, and green hues, corresponding to a respective decrease in collagen bundling. The data is presented as a ratio of the number of pixels of each color category over the total across all color categories per group. Data are mean ± SEM, n=4-7; *p<0.05 vs. Control Female.

To determine the alignment of smooth muscle fibers in the aorta, fiber angle was assessed by CurveAlign software. Representative images of the aorta from each group are shown on **Figure 6A**. Various additional ECM characteristics such as fiber straightness, length, and width were analyzed per animal in each group using CT-Fire software. HLU did not affect any of the studied parameters in either male or female groups (**Figure 6**). No differences were evident between sexes in control or HLU-treated groups (**Figure 6**).

**Figure 6.**
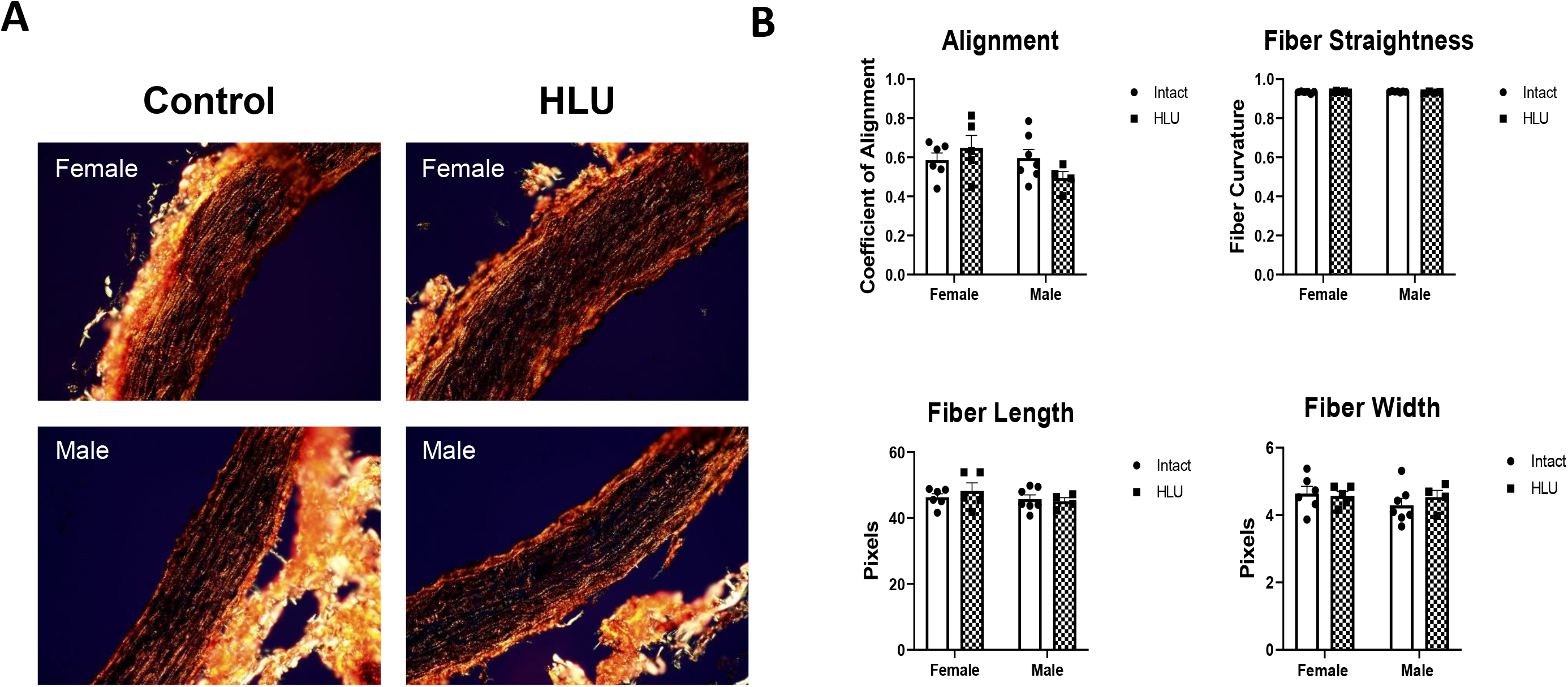
Collagen fiber characterization in female and male SD rats. (A) Female and male rat aortas in both control and HLU groups were collected after 14 days of either control or HLU exposure. Picrosirius red stained slides were imaged with polarized light, with representative images from each group shown. (B) ECM characteristics were analyzed using CT-FIRE software to assess the straightness, length, and width of individual fibers, with n=4-7 animals per group. Fiber alignment was quantified using CurveAlign software.

### 3.4 Changes in aortic G protein-coupled estrogen receptor in response to HLU

GPER staining in aortic media was lower in control males versus control females. HLU reduced the levels of GPER in females versus control females. No differences in aortic GPER were found in males in response to HLU (**Figure 7**).

**Figure 7.**
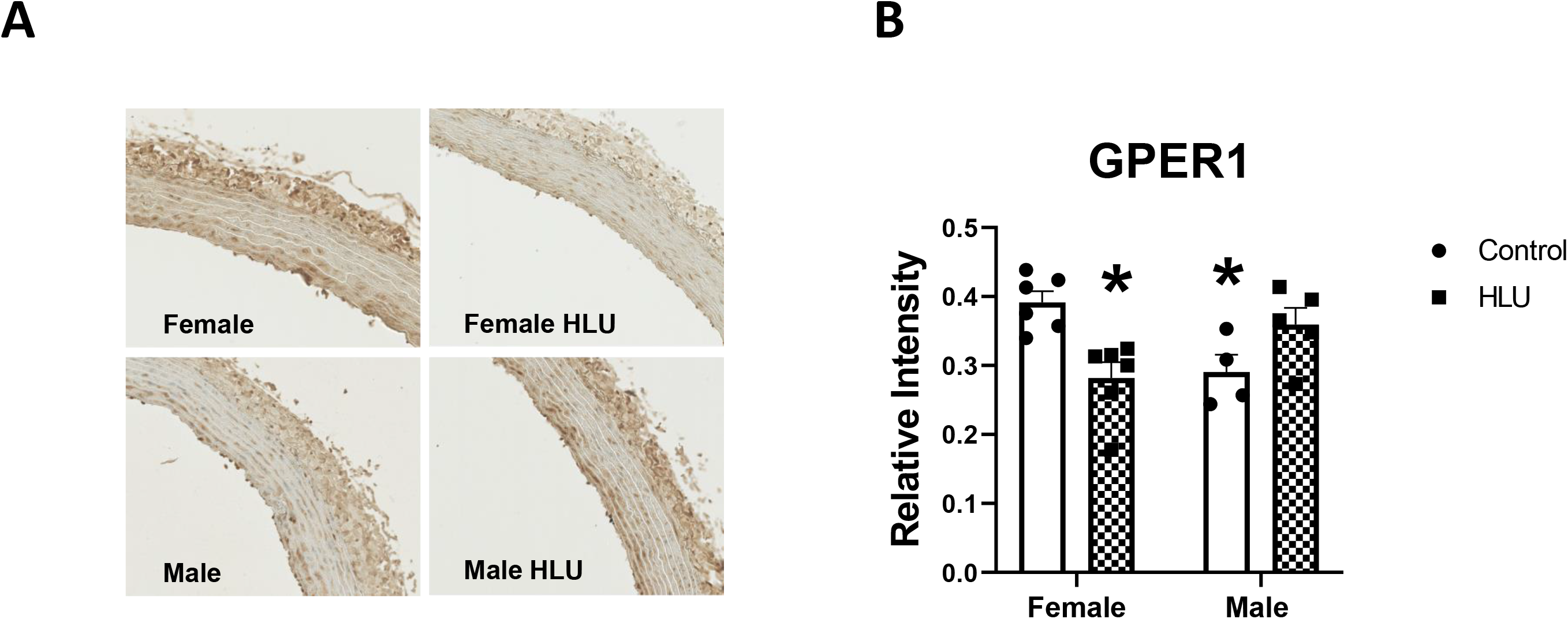
GPER1 levels in female and male SD rats in response to HLU. Clear bars indicate control unexposed animals; checkered bars indicate rats exposed to hindlimb unloading (HLU). Representative images (A) and data analysis are shown (B). Data are mean ± SEM, *p<0.05 vs. Control Female, n=4-6.

### 3.5 Inflammation and oxidative stress markers in aorta following the exposure to HLU

Male control rats had lower levels of PPAR-γ and COX-2 in the aortic media compared with control female rats (**Figure 8A, B)**. PPAR-γ levels decreased in female rats following HLU (**Figure 8A**). HLU did not affect the levels of COX-2, eNOS, or CD68 in the aortic media of female or male rats (**Figure 8B, C, D)**. However, p47 phox and 8-OHdG levels were increased in the aortic media in the female group after HLU compared with the female control group (**Figure 8E, F)**. 4-HNE was lower in control males versus control females, but no differences were found in the aortas of HLU-exposed females or males versus respective controls (**Figure 8G**).

**Figure 8.**
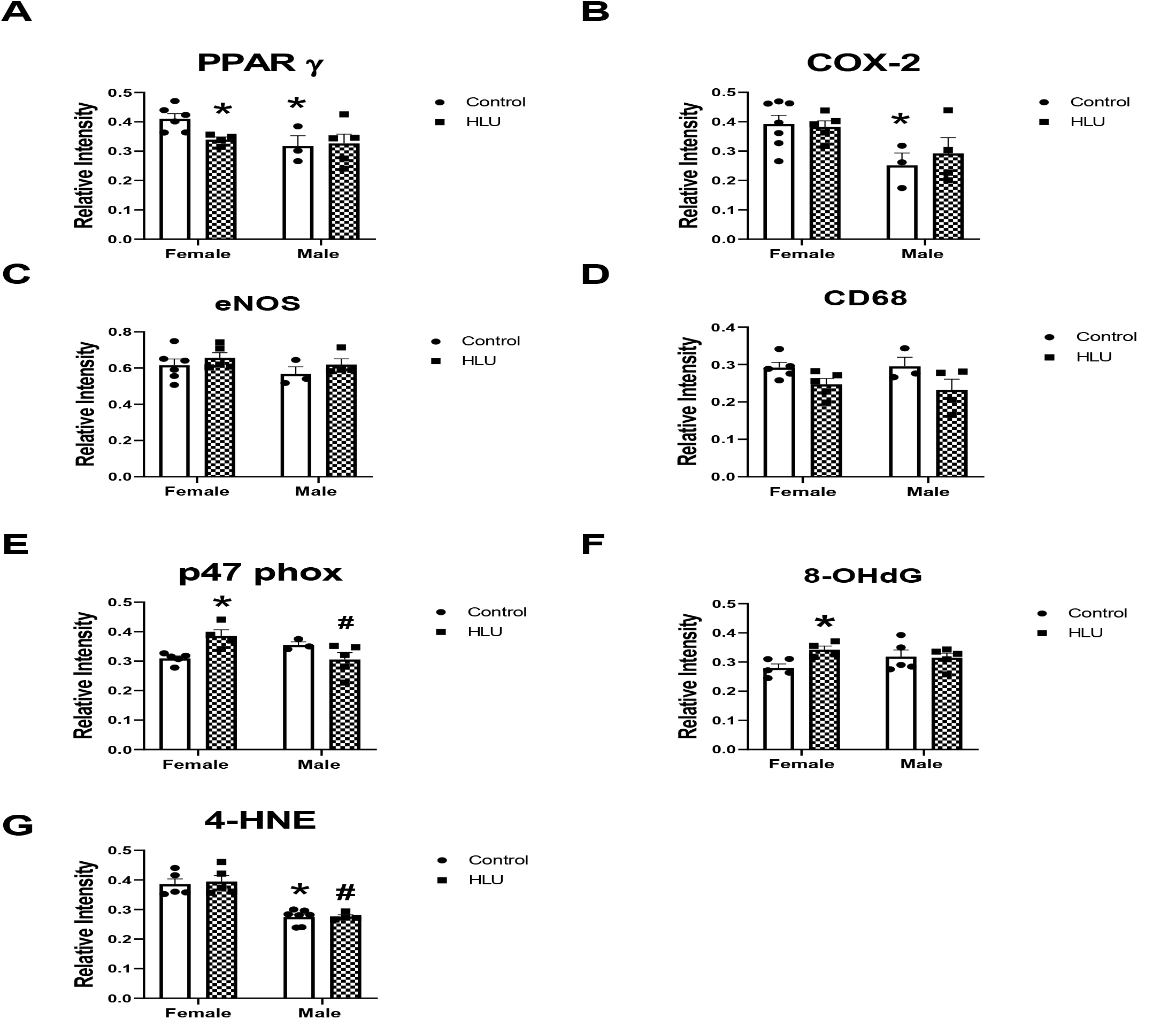
Aortic Injury markers in female and male SD rats in response to HLU. Data are mean ± SEM, *p<0.05 vs. Control Female; ^#^p<0.05 vs. HLU Female; ^^^p<0.05 vs. Male.

### 3.6 The effects of G1 on aortic PWV, injury markers, and uterine weight in female HLU SD rats

G1 administration reduced aortic arch PWV in female rats exposed to HLU (**Figure 9A**). No difference in uterine weight was evident in female rats after G1 treatment (**Figure 9B**). The levels of 8-OHdG levels decreased following G1 treatment in HLU female rats (**Figure 9C**). No differences in p47 phox or PPAR γ levels were detected in HLU females (**Figure 9D, E**).

**Figure 9.**
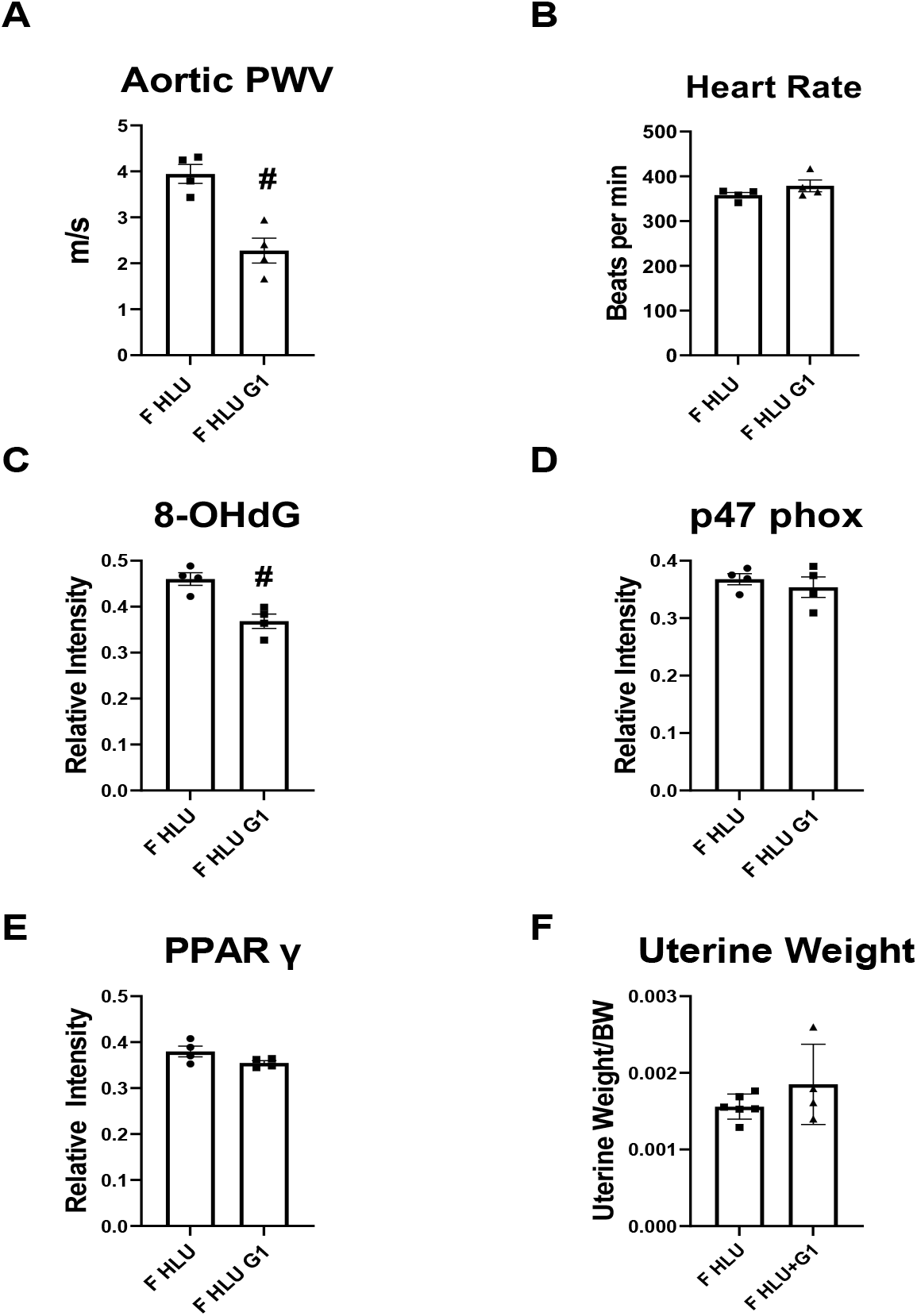
The response to GPER agonist in the female SD rats following HLU. Data are mean ± SEM, ^#^p<0.05 vs. HLU Female.

## 4 Discussion

Recent studies highlighted the incidence of arterial stiffness in astronauts following spaceflight. The exposure of rodents to ground-based analogs of microgravity via HLU also demonstrated development of arterial stiffness. However, few studies investigated sex differences and the potential sex-specific mechanisms leading to arterial stiffness due to microgravity. As space exploration continues to expand with longer and more distant missions, understanding the cardiovascular differences between female and male astronauts will be crucial to ensure the long-term health and personalized medical care for astronauts (1). In this study, we investigated sex-specific effects on the cardiovascular system in a rodent-based model of simulated microgravity via 14 days of exposure to HLU (17). The major findings of this study are that female and male SD rats respond differently to HLU demonstrating a greater arterial stiffness in females at the end of the exposure compared with males. Increased arterial stiffness was associated with lower levels of aortic GPER and PPARγ and increased oxidative stress markers (8-OHdG and p47 phox) in females but not males after HLU. The females also exhibited diastolic dysfunction after HLU. Furthermore, administration of G1, an agonist for the G protein-coupled estrogen receptor GPER, reversed the HLU-induced increase in PWV in female rats in association with lower 8-OHdG, suggesting that decreased GPER in the setting of simulated microgravity contributes to arterial stiffening. Our findings also suggest that the SD female rat is a suitable model to study mechanisms underlying the development of arterial stiffness in females in response to simulated microgravity.

It is well known that during spaceflight, the cardiovascular system adapts to the state of microgravity by a variety of changes including redistribution of blood flow with fluid shift towards the upper body and decrease in venous return and resistance. Sex differences were reported in the adaptations of the cardiovascular system to spaceflight, however only a few studies investigated the effects of microgravity on arterial stiffness by including subjects of both sexes, and even fewer have made explicit side-by-side comparisons between male and female subjects (1, 8, 10, 18-23). Although published reports also suggest that arterial stiffening is reversible in astronauts upon returning to Earth (24), the long-term consequences of the exposure to arterial stiffening are unknown but may influence future abnormal hemodynamic responses to stressors and the risk of developing cardiovascular disease (25–29) particularly in older females (30). A greater aortic PWV in the female SD rats compared to males after HLU suggests sex differences in the development of central arterial stiffening following the exposure to simulated microgravity in this model. PWV differs between women and men during various stages of life ranging from lower baseline PWV in females versus males pre-puberty, similar brachial-ankle PWV in both sexes during adulthood, and a greater increase in PWV in females after menopause suggesting hormonal influences on PWV (31–33). Furthermore, women undergoing cardiac stress often experience increased aortic wall thickness compared to men, highlighting a potential increased susceptibility of females to developing arterial stiffness (34). In contrast to other reports, our study did not find changes in arterial stiffness in male rats in response to HLU (22). The majority of these studies used Wistar rats or male C57Bl/6 mice (22), thus the response seen in SD rats may be different from those seen in other rodents. Furthermore, it has been reported that corticosterone levels are lower in resting male SD rats compared with Wistar rats (35), suggesting strain differences in the activity of hypothalamic-pituitary-adrenal (HPA) axis (36). Since hormonal regulation of vasculature underlies its response to stress, it is safe to assume that the response of male Wistar rats could be exacerbated by greater resting activities of HPA and corticosterone compared with male SD. Furthermore, the exposure length to HLU, age of rats, and the differences in strain responses to stress may explain the observed discrepancies. Thus, a longer exposure of SD males to HLU is potentially needed to develop the magnitude of change in PWV seen in Wistar rats.

Previous studies suggested that increased arterial stiffness due to microgravity may be a compensatory mechanism necessary to increase venous return and cardiac output after spaceflight (37). Although we have not directly measured total peripheral resistance, the increased stiffness in females was associated with maintained cardiac output and stroke volume after HLU. In addition, although there was a significant increase in ejection fraction in females after HLU, the ejection fraction values were within the normal range for rats at this age. The significance of these findings needs further investigation by examining the presence of subclinical left ventricle systolic dysfunction using more advanced features of strain rate imaging (38). In contrast, we found a significant increase in E/E’ ratio in female rats after HLU indicating a diastolic dysfunction. These data are in agreement with recent studies including the Framingham Heart Study that reported correlations of arterial stiffness with left ventricular diastolic function (39–41). It is possible that increased aortic stiffness leads to left ventricular hypertrophy followed by lower filling pressure and reduced coronary flow (41). Thus, the increased E/E’ ratio in females after HLU in addition to greater left common carotid artery stiffness in our study may indirectly suggest the reduction of left ventricle filling pressure. Arterial stiffness, pulse pressure amplification, and augmentation index were greater in women versus men in the Cardiovascular Abnormalities and Brain Lesions study, where the authors suggested that higher arterial stiffness may contribute to higher susceptibility of women to develop heart failure with normal ejection fraction (42). The return of cardiovascular system to normal functioning after spaceflight may be delayed by lower vasoconstriction of peripheral arteries or lower capacity to elevate vascular resistance in those who developed arterial stiffness emphasizing the need for longitudinal assessment of sex-specific risk of cardiovascular events in astronauts with arterial stiffness following long-term spaceflights (37, 43). In addition, other hemodynamic variables such as blood pressure can influence the extent of arterial stiffness. Higher blood pressures were reported in male SD or male Wistar rats compared to controls after 14 days of HLU (44, 45). Therefore, it is possible that the response of blood pressure after HLU was different in male versus female SD rats contributing to differences in PWV. Our future studies will incorporate blood pressure measurements during HLU exposure to establish the interactions of estrogen receptors and arterial hemodynamics.

Changes in mechanical forces (shear stress and pressure) or ECM composition are important for the regulation of aortic vascular smooth muscle cell (VSMC) support of the arterial compliance and the response of VSMC to increases in blood volume or pressure (46). We investigated the alterations in ECM components as a potential contributor to increased PWV. Male rats had greater expression of aortic α smooth muscle actin and myosin, however no differences in these cytoskeletal elements were detected in either female or male rats after HLU. In contrast, lower expression of α smooth muscle actin with potential switch to synthetic phenotype was reported in cerebral arteries suggesting vessel-specific alterations in cytoskeleton in response to HLU (47).

An increase in collagen cross-linking and deposition and decrease in elastin content may also lead to increased aortic stiffness. Our findings reveal no differences in collagen content after 14 days of HLU. These data are consistent with reports demonstrating that structural remodeling may not be the primary mechanism leading to arterial stiffness in astronauts after spaceflights (24) suggesting that other factors such as systemic (neurohumoral) or local endothelial factors may have an impact on the development of arterial stiffness (10, 22). However, others reported that HLU resulted in increased deposition of extracellular collagen in the basilar artery but not femoral artery in rats (48). The hydroxyproline content as a measure of collagen content was increased in male Wistar rats, however collagen subtype composition was not different between control and HLU-exposed rats in this cohort (22). These studies suggest that hypertrophic response to HLU is vascular bed specific. Furthermore, our data revealed no differences in fiber characteristics including alignment, straightness, length, or width suggesting that 14-day exposure to HLU does not induces changes in collagen fiber characteristics in SD rats. This was a surprising finding considering a possibility of an increased pressure on collagen fibers and their gradual strengthening under pressure due to HLU (49). Increased stiffness of collagen fibers may also affect smooth muscle cell phenotype, cell proliferation, or cause downregulation of vascular contractility (50, 51). Increased fiber angle may also indicate greater alignment between smooth muscle fibers potentially contributing to increased stiffness (52). Furthermore, highly dense and stiff matrices can promote crosstalk between estrogen receptors and prolactin supporting the hypothesis that changes of the physical environment, such as in microgravity, can be affected by estrogen receptors (53). Unlike collagen, another major ECM protein, elastin, is responsible for elasticity, reversible expansion and relaxation of large arteries (54). Despite elastin being a critical component of the aorta’s ability to deform under stress, few studies have investigated elastin in response to HLU exposure. No statistically relevant differences in elastin expression or collagen-to-elastin ratio were found in either female or male rats after HLU exposure. However, the female HLU group had a wide standard deviation which consisted of several subjects having decreased elastin expression after HLU exposure compared to the mean. For those individuals, the cause of decreased elastin after HLU exposure is unclear. Elastin in the aorta could decrease with age or in the setting of arterial stiffness. However, similar to our study, aortic elastin content was not altered in male Wistar rats after HLU (22).

Sex differences in the cardiovascular system may depend in part on the levels of sex hormone receptors and signaling induced by their actions. In non-pregnant female mice, aortic PWV was reported to be lower in estrus cycle compared with diestrus suggesting that the development of vascular stiffening may in part be influenced by the levels of ovarian hormones (55). Indeed, our preliminary data agrees with previous reports suggesting that the removal of sex hormones via ovariectomy increases arterial stiffening in the female rats. Furthermore, our preliminary studies show that HLU did not change PWV in aorta or carotid artery of ovariectomized rats (Figure S4). Similarly, ovariectomy did not exacerbate muscle loss in Fisher female rats during the exposure to either simulated microgravity or partial-gravity environments (56). These data together with our findings suggest that the activation of local vascular estrogen receptors may have a greater impact on the development of arterial stiffness after HLU than changes in the ovarian hormones. Furthermore, arterial stiffness is associated with genetic variants of estrogen receptor subtypes suggesting a link between arterial stiffness and estrogen receptors expression (57). Although the data on the status of estrogen receptors in response to spaceflight or microgravity analogs are limited, Holets et al. reported a downregulation of estrogen receptor α mRNA and protein in the uterus and ovary of mice after the spaceflight (58). Importantly, recent studies highlighted the role of estrogen receptors and GPER in arterial stiffness (59). GPER is a membrane-bound estrogen receptor that induces vasodilation and reduces blood pressure (60). GPER activation via its analogs such as G1 ameliorates aortic remodeling and reduces arterial stiffness in ovariectomized and hypertensive rats (59). Our data demonstrate for the first time that HLU exposure for 14 days reduces the levels of GPER in the aortic intima of females without a similar effect in males.

Our data also show lower PPARγ levels in aorta of female rats in response to HLU (61). PPARγ is a member of the nuclear receptor superfamily involved in estrogen-based protection of the cardiovascular and renal systems (62, 63). PPARγ is also closely linked to cardiovascular and metabolic dysfunction and therefore has the potential for being targeted by HLU which induces cardiovascular changes similar to aging or menopause (64). Considering the role of PPARγ in the regulation of vasodilation, vascular redox state, bioavailability of nitric oxide, as well as regulation of VSMC stiffness (65), our data implies that lower PPARγ may result in reduced protection against arterial stiffness caused by the exposure to HLU. Furthermore, lower PPARγ in the aorta of female rats was associated with lower GPER levels suggesting the interaction between GPER and PPARγ in the setting of HLU. Both GPER and PPARγ can reduce oxidative stress and pro-inflammatory stimuli in vasculature. Our data revealed increased levels of DNA damage marker 8-OHdG in the aorta of HLU females suggesting increased oxidative stress. Furthermore, the absence of differences in the levels of lipid peroxidation marker 4-HNE suggests a more pronounced DNA damage (66). In agreement, a 72-hour exposure to simulated microgravity through a rotary cell culture system resulted in increased expression of the DNA damage markers in human promyelocytic leukemia cells (67). Previous reports demonstrated increased circulating levels of 8-OHdG in rats after HLU, in human patients with cardiovascular disease, as well as in patients with diabetes where higher 8-OHdG was positively correlated with PWV (68–70). Lower levels of antioxidant enzymes catalase and glutathione peroxidase were reported in skeletal muscle of male SD rats after 28 days of HLU suggesting that lower activity of antioxidant enzymes may also contribute to the increase of superoxide levels after HLU (71). In addition, oxidative stress may increase vascular inflammation or impair the balance of vasodilator and vasoconstrictor mediators, therefore we investigated the expression pattern of CD68-positive macrophage/monocytes, eNOS and COX-2 in the aorta after HLU. The eNOS regulation in the vasculature in response to HLU may depend on the vascular bed or the length of the exposure (43, 72). Lower nitric oxide availability due to formation of peroxynitrite has been suggested as a contributing factor to superoxide production in vasculature after HLU (73). However, our data revealed no differences in eNOS or COX-2 proteins among studied groups. Future studies will assess nitric oxide bioavailability and aortic vascular reactivity in response to nitric oxide and COX-2 inhibition to determine if functional vasodilatory responses are altered in female rats after HLU.

Finally, to determine whether lower aortic GPER contributes to the development of arterial stiffness in response to HLU, female rats were treated with G1, a selective agonist of GPER. Our data show that GPER agonist G1 reversed increased arterial stiffness in the female rats, changes associated with the reduction in oxidative stress suggesting that lower GPER may contribute to the development of arterial stiffness after HLU, and the protective actions of GPER may involve reduction of vascular oxidative stress (60).

### Summary

Simulated microgravity as modeled by HLU results in changes in the cardiovascular system. The present study reports varied responses of cardiovascular system in female and male rodents when exposed to HLU. Although the cellular mechanisms of the interaction between microgravity, the cardiovascular system and estrogen receptors in response to HLU need further investigation, our study reveals that lower GPER in the females contributes to the development of arterial stiffness during HLU and that the SD rat is a suitable model to study these interactions.

## Abbreviations

SD: Sprague Dawley rat
HLU: Hindlimb unloading
PWV: pulse wave velocity
GPER: G protein-coupled estrogen receptor
PPARγ: peroxisome proliferator-activated receptor γ
OHdG: 8-hydroxy-2’-deoxyguanosine
SANS: spaceflight associated neuro-ocular syndrome
ROIs: regions of interest
4-HNE: 4-hydroxynonenal
COX-2: yclooxygenase-2
eNOS: endothelial nitric oxide synthase
HPA: hypothalamic-pituitary-adrenal axis
VSMC: vascular smooth muscle cell
ECM: extracellular matrix

## Author Contributions

Ebrahim Elsangeedy performed the experiments, acquired and interpreted the data; Dina N. Yamaleyeva analyzed and interpreted the data, wrote and revised the manuscript; Nicholas P. Edenhoffer analyzed, acquired and interpreted the data, revised the manuscript; Allyson Deak, Anna Soloshenko, Omar Shaltout, and Nildris Cruz Diaz analyzed, acquired and interpreted the data; Jonathan Ray and Xuming Sun performed the experiments, analyzed the data, revised the manuscript; Brian Westwood analyzed and interpreted the data; Daniel Kim-Shapiro, Debra Diz, Shay Soker, and Victor M. Pulgar interpreted the data, revised the manuscript interpreted the data, revised the manuscript; April Ronca consulted on study design, interpreted the data, revised the manuscript; Jeffrey S. Willey provided expertise on HLU model and study design; Liliya M. Yamaleyeva conceived and designed the research, analyzed and interpreted the data, wrote and revised the manuscript.

## Acknowledgements

These studies were supported by the WFUSM Cardiovascular Sciences Center Pilot Award to LMY, R. Odell Farley Hudson Research Fund, and the Hypertension and Vascular Research Center. We would like to acknowledge the Preclinical Ultrasound and Photoacoustic Imaging Core of Wake Forest University School of Medicine (WFUSM) supported in part by the Wake Forest Clinical and Translational Science Institute (NIH NCATS UL1TR001420). We also acknowledge the editorial assistance of Shea Gilliam, PhD.

## Data Availability Statement

The data that support the findings of this study are available in the Materials and Methods, Results, and/or Supplemental Material of this article.

