## Supplementary Figures for "Sex-Specific Cardiovascular Adaptations to Simulated Microgravity in Sprague-Dawley Rats"

### Supplementary Material

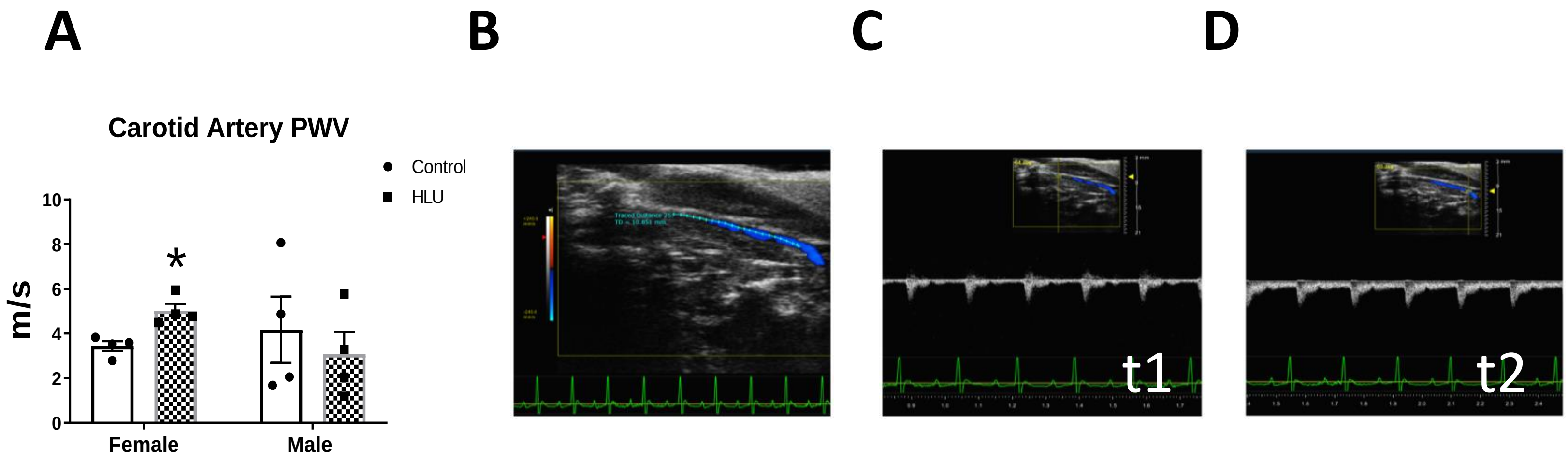


**Supplementary Figure S1.** *Carotid artery pulse wave velocity (PWV) in female and male rats in response to HLU*. Clear bars indicate control unexposed animals; checkered bars indicate rats exposed to hindlimb unloading (HLU) **(A)**. PWV was calculated as PWV=D/pulse transit time Δt, where D is the distance between two points in mm (t2 and t1) of carotid artery and t is the signal transit time estimated between R on electrocardiogram and the beginning of the Doppler wave in ms **(B-D)**. Data are mean ± SEM, *p<0.05 vs. Control Female.

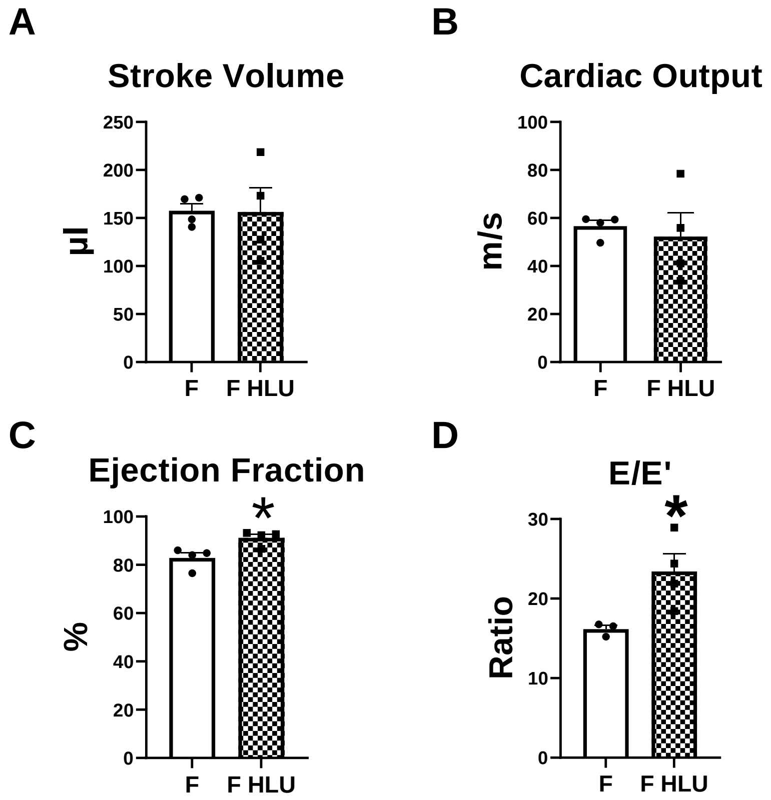


**Supplementary Figure S2.** *Cardiac function characterization in female SD rats after 14 days of HLU.* Data are mean ± SEM, *p<0.05, vs. Control Female (F), n=4 per group.

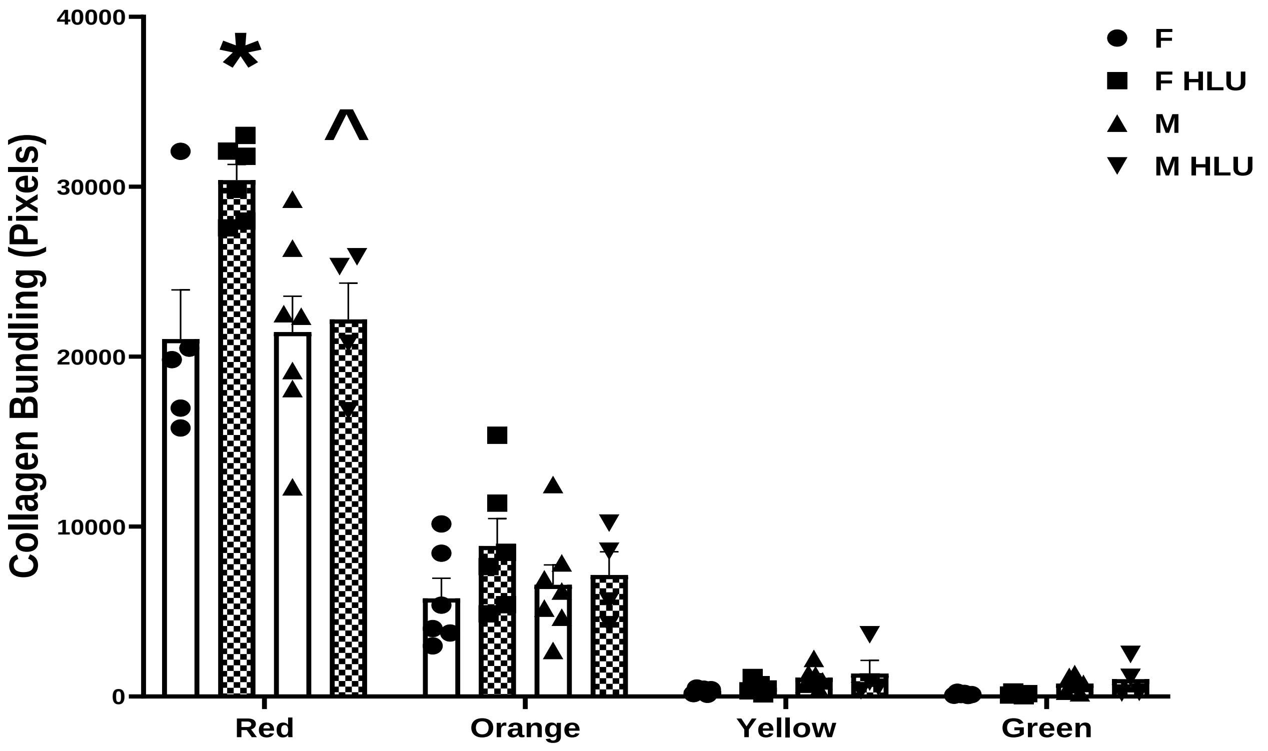


**Supplementary Figure S3.** *Collagen fiber bundling in the aorta of female and male SD rats in response to HLU*. Clear bars indicate control animals; checkered bars indicate rats exposed to hindlimb unloading (HLU). Picrosirius red stained slides were imaged with polarized light, and the pixel values of representative regions were binned into red, orange, yellow, and green hues, corresponding to a respective decrease in collagen bundling. Data are mean ± SEM, *p<0.05 vs. Control Female; n=4-7 rats per group.

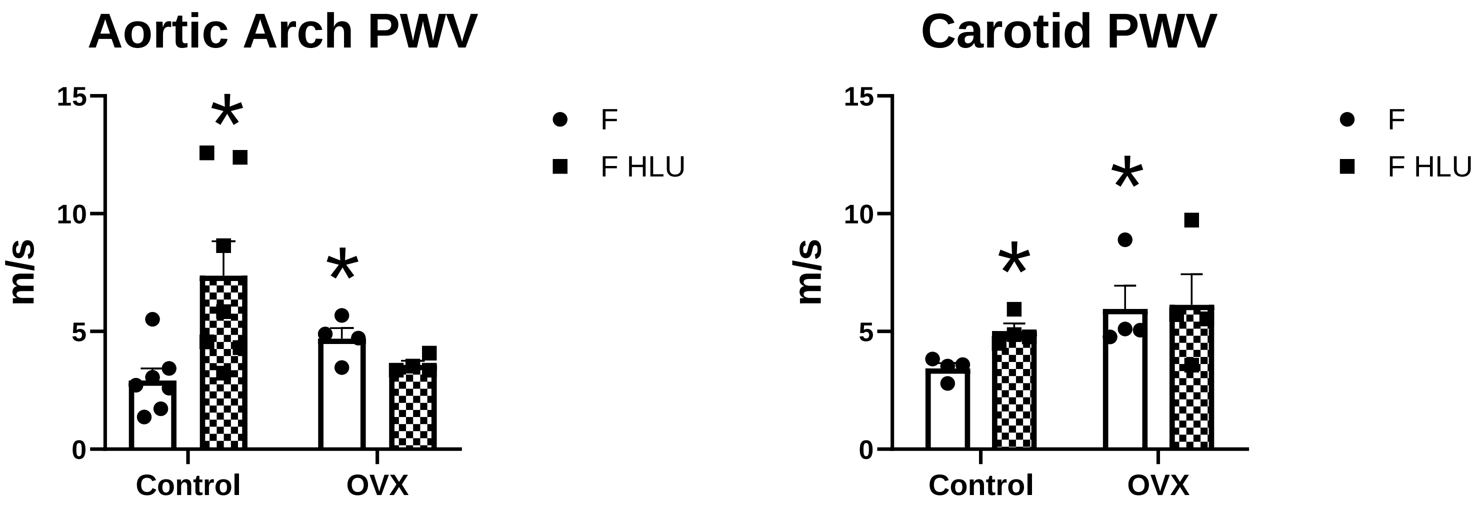


**Supplementary Figure S4.** *The effect of ovariectomy on aortic arch and carotid artery pulse-wave velocity (PWV) in female SD rats after 14 days of HLU.* Data are mean ± SEM, $^{*}p<0.05$, vs. Control Female (F), n=4 per group.
